# Constructing Causal Cytokine Networks for Autoimmune and Inflammatory Diseases Target Identification

**DOI:** 10.1101/2022.01.30.478394

**Authors:** Yanqing Chen

## Abstract

Cytokines are cell signaling proteins that act as inducers of physiological responses including the activation and differentiation of innate and adaptive immune cell types. In some cases, this mediates the initiation or progression of inflammatory and auto-immune diseases such as septic shock, rheumatoid arthritis, psoriasis, and inflammatory bowel disease. Many cytokines are pleiotropic, meaning that they can have effects on multiple cell types and play several distinct functions. On the other hand, multiple cytokines might have similar function and be redundant or degenerate. Some cytokines also regulate the expression and functions of other cytokines. Such complex cause-effect relationships can be modeled computationally by building “cytokine networks”. Those networks describe intricate interactions between different cytokines in their associated tissue environment and cell types. Here we propose a computational approach to find sensitive and “causal” nodes within a cytokine interaction network by identifying whether persistent positive feedback (self-loop) exists on that cytokine in its mRNA transcription network. Based upon two microarray gene expression datasets from immune stimulation experiments using up to 15 different cytokines, we identified TNFα as a “causal” cytokine in the colonic epithelial cell line HCT8, and IL-23, IL-1β and TGFβ as “causal” cytokines in primary myofibroblasts. We hypothesize that a cytokine with a persistent positive feedback loop that amplifies its transcription and secretion could be identified as “causal” for the transcriptional regulation of other cytokines and potentially, downstream disease phenotypes. The existence of such positive feedback loop may determine whether targeting suppression of such cytokine will result in clinical efficacy.

## I. Introduction

Targeting cytokines is an important therapeutic strategy in several immune related diseases. Among them, anti-TNFα antibodies have shown efficacy in several auto-immune diseases such as Crohn’s disease, ulcerative colitis, rheumatoid arthritis, psoriasis, etc. In contrast, antibodies targeting other cytokines such as IL-17A have shown efficacy in a subset of auto-immune diseases such as psoriasis (Farahnik et. al., 2016). The success of TNFα inhibitory therapy on several immune-related diseases suggests it is an essential and important node in the “fragile communication networks of cytokines” (Schett et al. 2013). Determining a cytokine’s function and “causal” role for immune-related disease phenotypes could increase our understanding of disease mechanisms and help finding other valuable targets for specific diseases. In many immune-related disease phenotypes, protein or gene expression levels of TNFα are increased as compared to healthy controls. Yet, simple elevation of protein/gene expression level in a disease state does not mean the gene/protein is causal for the disease nor targeting it would be effective for therapeutic efficacy. The increase in expression might only suggest it is correlated with disease progression. Differentiating “causality” from “correlation” is a crucial step in identifying a promising disease target.

Here we hypothesize that TNFα is in a “positive-self-regulated” state in disease-relevant tissue types or responder patient subpopulations, and that state can be used to discriminate “causal” vs. “reactive” nodes in a protein/cytokine network: a persistently generated cytokine exhibiting a positive feedback loop would more likely drive the disease phenotype. This “positive-self-regulated” state means, that within a relative broad range of TNFα concentration levels, there is a positive feedback loop between TNFα level and its transcription/production. This positive feedback loop is stronger than suppressive controlling factors from other genes/proteins, such that it controls (“causal for”) other genes/phenotypes rather than the other way around. In comparison, if its self-positive feedback is weak, or dominated by negative self-feedback (controlled by other factors), a gene/protein/cytokine can be viewed as being “reactive”. “Reactive” genes/proteins will have a tighter (smaller) range of homeostatic values or follow other proteins changes. Perturbing them would have less of a down-stream effect.

The phenomenon of feedback control on a cytokine is related to the concept of “autocrine” and “paracrine” in immunology, yet it is more general because of emphasizing the dynamics of cytokine interactions over time rather than cell type or physical location. Autocrine, by definition, means a cytokine is released and acting back on the cell which produces it, while paracrine means the cytokine is acting on other nearby cells or cell types (Caldwell et a., 2014). In a sense, these two concepts represent a network-type of cytokine action based upon cellular proximity. Through identifying self-feedback loops, our method here could potentially detect autocrine behavior if experiments are done in a single cell-type. In addition, from molecular profiling of samples with multiple cell types we could potentially detect more complex feedback interactions involving autocrine and paracrine communication among multiple cell types and locations.

In this article we will describe some of the mathematical background for the Causal Cytokine Network concept and propose ways to identify dynamic “causal” and “reactive” nodes in a cytokine network, then list some preliminary data analyses and published literatures that support this idea. This method is built upon the general concept of “cytokine network” (Morel et al., 2017) and provides a practical means of identifying directional interactions in complex biological networks.

## II. Method: Apply concepts of dynamical causal networks and homeostatic control

The immune system responds to stimuli by initiating cellular and molecular changes after encountering challenging endogenous or exogenous signals. That leads to temporal changes in the expression of relevant proteins/cytokine genes, from hours to days, as measured by their mRNA levels and protein concentrations. Such process can be modeled by dynamic system equations (i.e., Ordinary Differential Equations – ODEs, see for example Waito et al. 2016). Basically, ODEs describe the rate of change (*dx/dt*) of a specific protein/cytokine as a function of the current concentration or gene expression level of its own, plus any other genes/proteins that might influence (i.e., control) its rate of change.

Figure 1 describes a simplified network of three dynamic variables (*x, y, z*) which could model the temporal concentration changes of three cytokines, for example, TNFα, IL-1β and IL-10. Equation (1) is a simple form to describe a positive self-regulated feedback variable (in fact, an unstable one), only positively controlled by itself. As a result, any slight increase of x will lead to exponentially larger increases of *x* at a future time point, resembling the rapid uncontrollable increases of TNFα levels in disease states such as septic shock or “cytokine storm” (a more detailed mathematical modeling of cytokine storm using ODEs is presented in Yiu et. al. 2012). Please noted that this is a simplified positive feedback equation. In reality, biological positive feedback might only exist within a physiological range, which could be more appropriately modeled by a modified version of Equation (1), such as *dx/dt=f(x)* where the function *f(x) >0* when x is between *x_min_* and *x_max_*, and *f(x)<0* when *x* is outside of that range.

**Figure 1.**
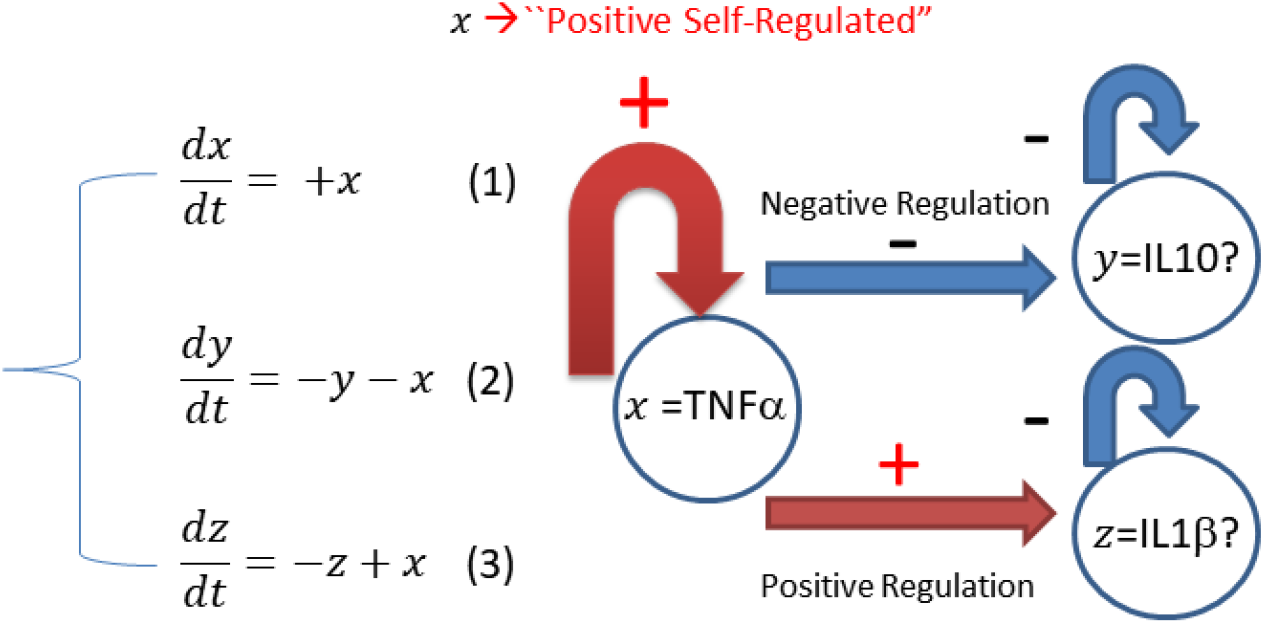
A three-variable system of Ordinary Differential Equations (ODEs) describes simplified positive (TNFα) and negative self-regulated (IL10 and IL-1β) dynamic interactions between three cytokines in a mathematical form. This schematic is used as a basis to model the cytokine network derived from in vitro stimulation data (see main text).

Equation (1) is an example of a strong self-regulated positive variable that not controlled by any other variables. In comparison, Equations (2) and (3) describe variables *y* and *z* as self-regulated negative variables that at the same time controlled by variable *x*. Since they both are negatively self-regulated, they will not respond to self-stimulation, but their values will follow *x*. For example, IL-1β increases when TNFα increases, and IL-10 might decrease if TNFα has an inhibitory effect on IL-10’s anti-inflammatory properties. The above 3-variable model can be used as a basis to construct dynamic causal networks, where *x* is a causal node controlling two downstream variables *y* and *z*. Their differences in terms of “causal” vs. “reactive” can also manifest in their homeostatic values, which in the equations can be determined when *dx/dt*, *dy/dt* and *dz/dt* are set to 0. As described above, homeostatic values of x will cover a wider range and be more sensitive to external input or perturbations, while *y* and *z* will be more stable as long as *x* is not changed.

## III. Experiment: Identify causal cytokines and self-regulated positive feedback loop via*in vitro* stimulation of colonic epithelial cells and myofibroblasts

A major difference between “causal” vs. “reactive” in the above dynamic network is whether homeostatic values are sensitive to perturbations, more specifically, whether a change in the variable’s own value will be stabilized or amplified. For many cytokines, such distinction can be determined via studies that monitor the effects of self-stimulation, i.e., whether a cytokine’s concentration or expression level will respond (increase or decrease) to its own production. We designed two sets of *in vitro* experiments to test effect of cytokine stimulation (induction) on two types of cells that are relevant to Inflammatory Bowel Disease (IBD), the human colonic epithelial cell line HCT8 and human primary intestinal myofibroblasts. These two types of cells (commercially purchased from ATCC) were stimulated with up to 13 cytokines in at different concentrations in the culture media. Some of the cytokines were also added in combination. In the following section we demonstrate how to construct cytokine networks from these cell-based stimulation studies, and how to identify potential “causal” and “reactive” cytokines from the inferred network.

In the first experiment, whole genome gene expression was profiled using Affymetrix microarrays in 168 HCT8 cell culture samples under various cytokine conditions for 20 hours. In total, 13 immune stimulatory conditions, each with three doses and technical triplicates for each dose, plus 1 group of combination stimulatory conditions (IL-22 + IL-1β; IL-22 + TNFα; IL-22 + TNFα + IL-1β) were analyzed, together with 42 control samples without any stimulation (3 replicates of control samples done for each stimulatory condition to minimize plate or batch effects). The second set of cytokine stimulation was done using intestinal myofibroblast primary cells. They were stimulated with cytokines or TLR ligands (PolyIC, Pam3CSK4, LPS, IL-33, IL-22, IL-17A, IL-17E, IL-15, TNFα, IL-23, TGFβ, IL-1β, IL-12, IL-4 and IL-13) along with control (non-stimulated) cells. Microarray data set profiled from these experiments were imported into ArrayStudio software (Qiagen OmicSoft) for statistical analysis. Cytokine stimulation signatures (i.e., ratio of stimulated samples versus their corresponding non-stimulated control samples) were generated for each stimulation condition using General Linear Model (GLM) with stimulation condition as main factor to compare. Genes that were significantly different under the cytokine stimulation versus non-stimulation control by passing a False Discovery Rate (FDR) <0.05 threshold with larger than 1.5fold changes were considered as stimulation signatures in each cell type.

## IV. Results

Figure 2 was generated from gene expression profiles of TNFα and IL-23 in the colon epithelial cell line (HCT8) under various cytokine stimulation conditions at 3 different doses for each condition. The top variable plot on TNFα gene expression levels showed it to be upregulated by self-stimulation, and slightly upregulated by IL-1β, and IL-1β + IL-22 or IL-1β + TNFα) combined stimulation. Figure 2(b) shows that IL-23 gene expression is not responsive to self-stimulation, but partially upregulated by TNFα and combined stimulation of that cytokine with IL-1β and IL-22. From these two gene expression profiles, we can construct the “cytokine interaction network” shown in the middle of Figure 2, where TNFα is found to be positively self-regulated, and potentially causal for the induction of IL-23 expression. It is worth pointing-out that TNFα is somewhat responsive to IL-1β and IL-1β + IL-22 in combination, yet neither IL-1β nor IL-22 self-stimulation their production or that of any other cytokine tested (data not shown). This would suggest that TNFα is sensitive to IL-1β and IL-22, with HCT8 cells unlikely to be the source for their production. In this sense TNFα is a “causal” cytokine in this specific cell type, but not IL-1β, IL-22 or IL-23. It is very likely that TNFα is causal for induction of many other cytokines, IL-23 being just one example detected by our analysis. Scanning of the “cytokinome” on microarray datasets would reveal the complete network.

**Figure 2.**
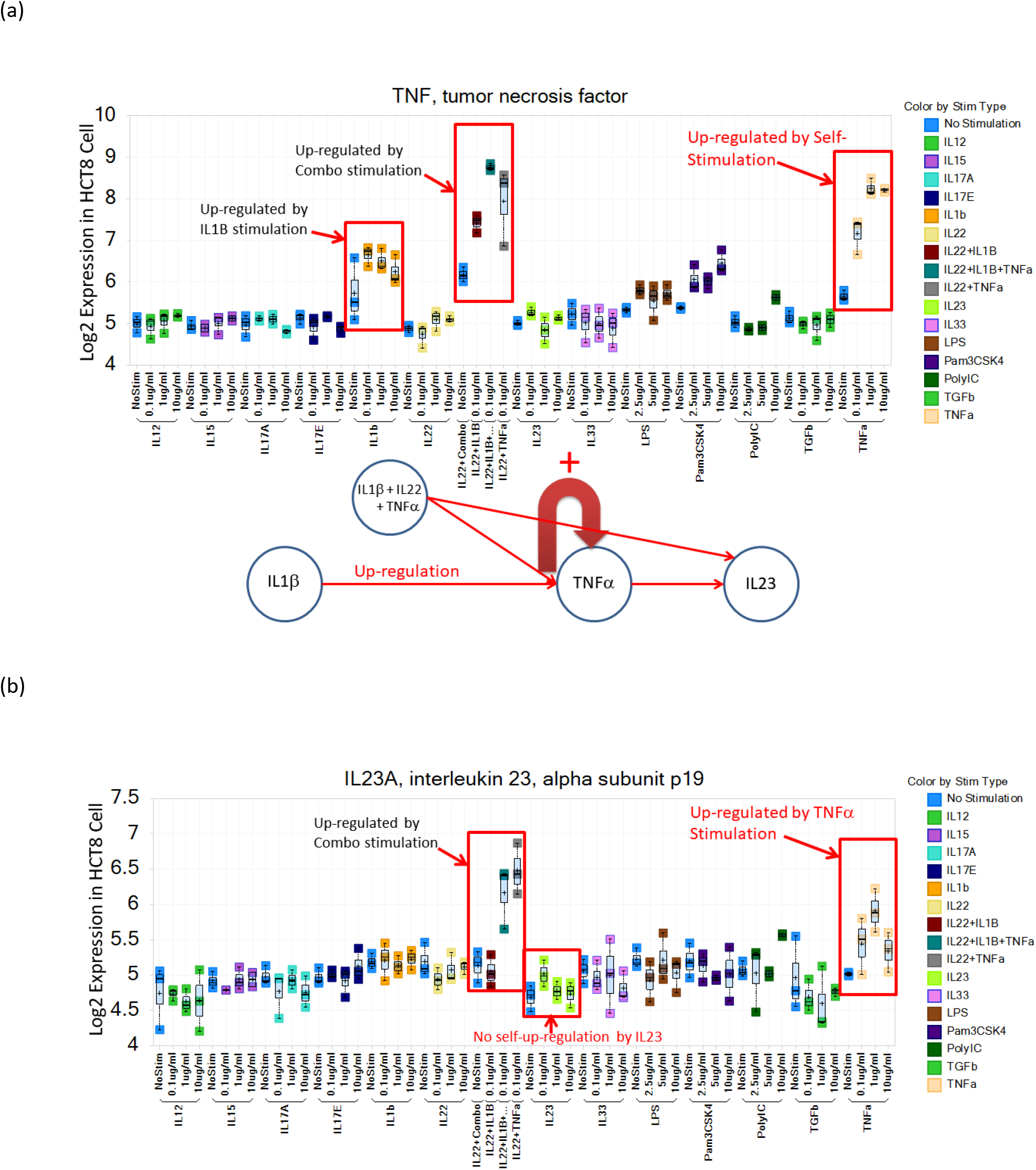
Gene expression profiles from two genes TNFα and IL23 under 13 stimulation conditions in the HCT8 cell lines. Top and bottom panels are Affymetrix microarray expression intensity values (in log2 scale) for TNFα (a) and IL23 (b), each column plotted in the panel are from 3 repeats of one type of stimulation at one specific dose labeled below, color of the dots represents the type of stimulation listed on the right bar plot where blue color represents the non-stimulated control. Stimulation ratios were calculated as the average difference between the stimulation plus dose condition versus the nearest control samples which are from the same cell culture plate without any stimulation applied. From both panels we can see that when TNFα or IL23 do not respond to a specific stimulation (for example, under IL12), their gene expression levels are about the same as non-stimulated control (around 5 in log2 scale for TNFα and IL23). If they respond, their expression values are significantly different from the non-stimulated control samples in blue as circled in the 5 specific stimulation conditions (including one self-stimulation for TNFα). The network plot between two panels summarizes the stimulation responses in a cytokine network form.

Figure 3 summarizes the overall cytokine interactions under the 13 stimulation conditions in a network graph using the open source software Cytoscape. It shows the mRNA expression regulation network based upon the cytokine stimulation experiment including only the cytokine nodes (interleukins, TNFα and TGFβ) with > 1.5fold changes under FDR (False Discovery Rate) <0.05 for any of the 3 doses tested from colon epithelial cell line (HCT8) stimulation experiment, where only TNFα is up-regulated by self-stimulation and marked in red (solid-line means up-regulation and dashed-line means down-regulation). As described before, the TNFα self-loop suggests that TNFα is a causal node in the cytokine network in the HCT8 cell type because it has the ability to amplify its own gene expression and to control other genes.

**Figure 3.**
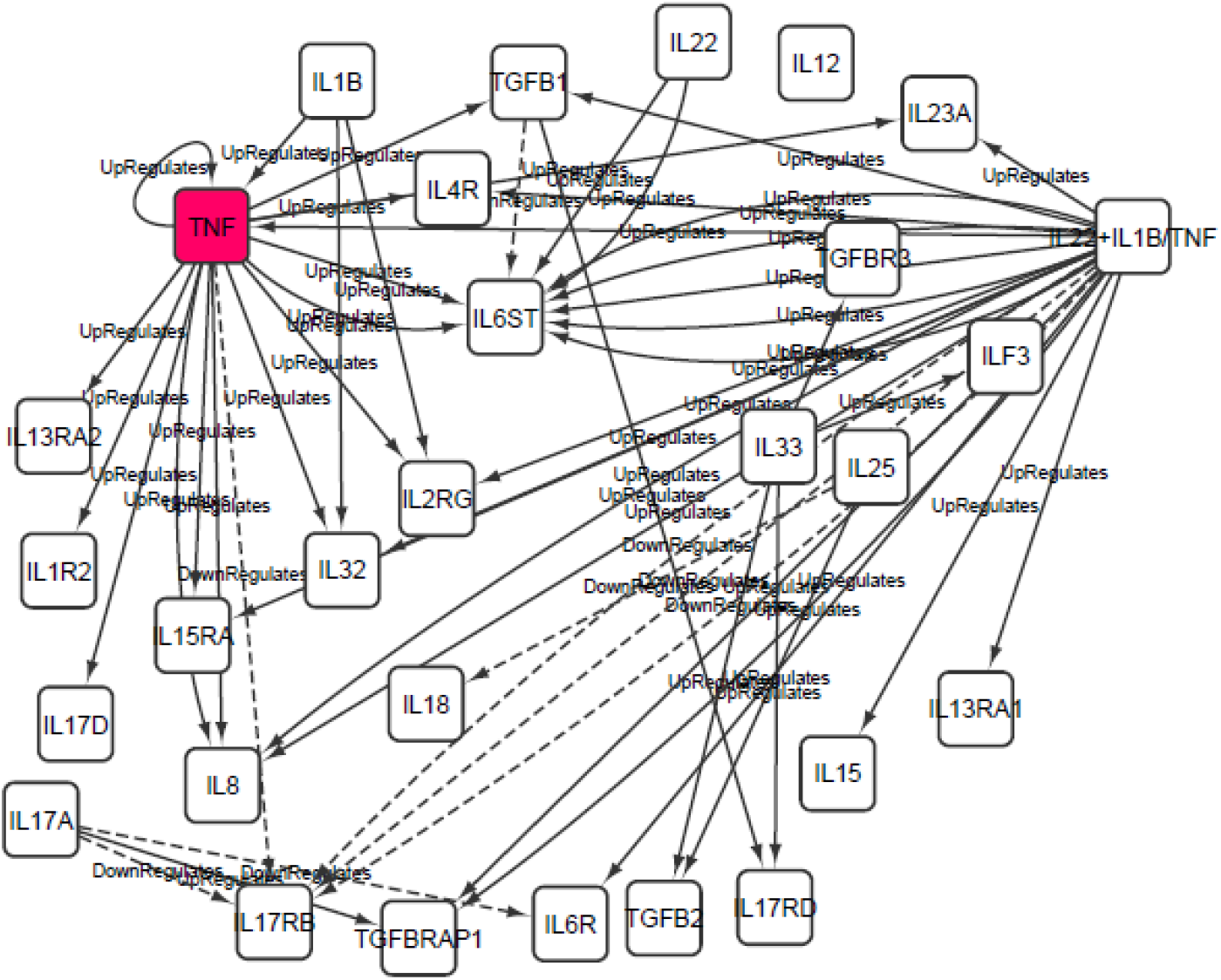
Cytokine network based upon the stimulation experiment from colon epithelial cell line HCT8. Directed lines represent the stimulation where the originating node is the cytokine stimulation, the end of an arrow represents other cytokines significantly modulated by the stimulation conditions in at least one dose tested (with > 1.5fold changes under FDR - False Discovery Rate <0.05). Solid lines represent up-regulation and increasing expression of target cytokines, and dashed lines represent down-regulation. Notice that only TNFα has a self-loop in this network which is emphasized as a red box here. This suggests that in the colon epithelial cell lines, only TNFα amplify its own gene expression out of the 13 cytokine stimulations tested.

Cytokine interactions might depend upon cell, tissue type and disease state. Figure 4 shows the gene expression profiles of IL-1β and TNFα based upon the second set of cytokine stimulation experiment performed on intestinal myofibroblast primary cells under similar experimental conditions. It demonstrates the interaction network between TNFα, IL-1β and IL-23. Besides TNFα, IL-1β and IL-23 also show some degrees of positive self-regulations, with interactions showing different directionality. This suggests that “causal” vs. “reactive” cytokine interactions are dependent upon tissue type and the specific context. This should not be surprising, as interactions between cytokines are likely be contextual in different cell and tissue types.

**Figure 4.**
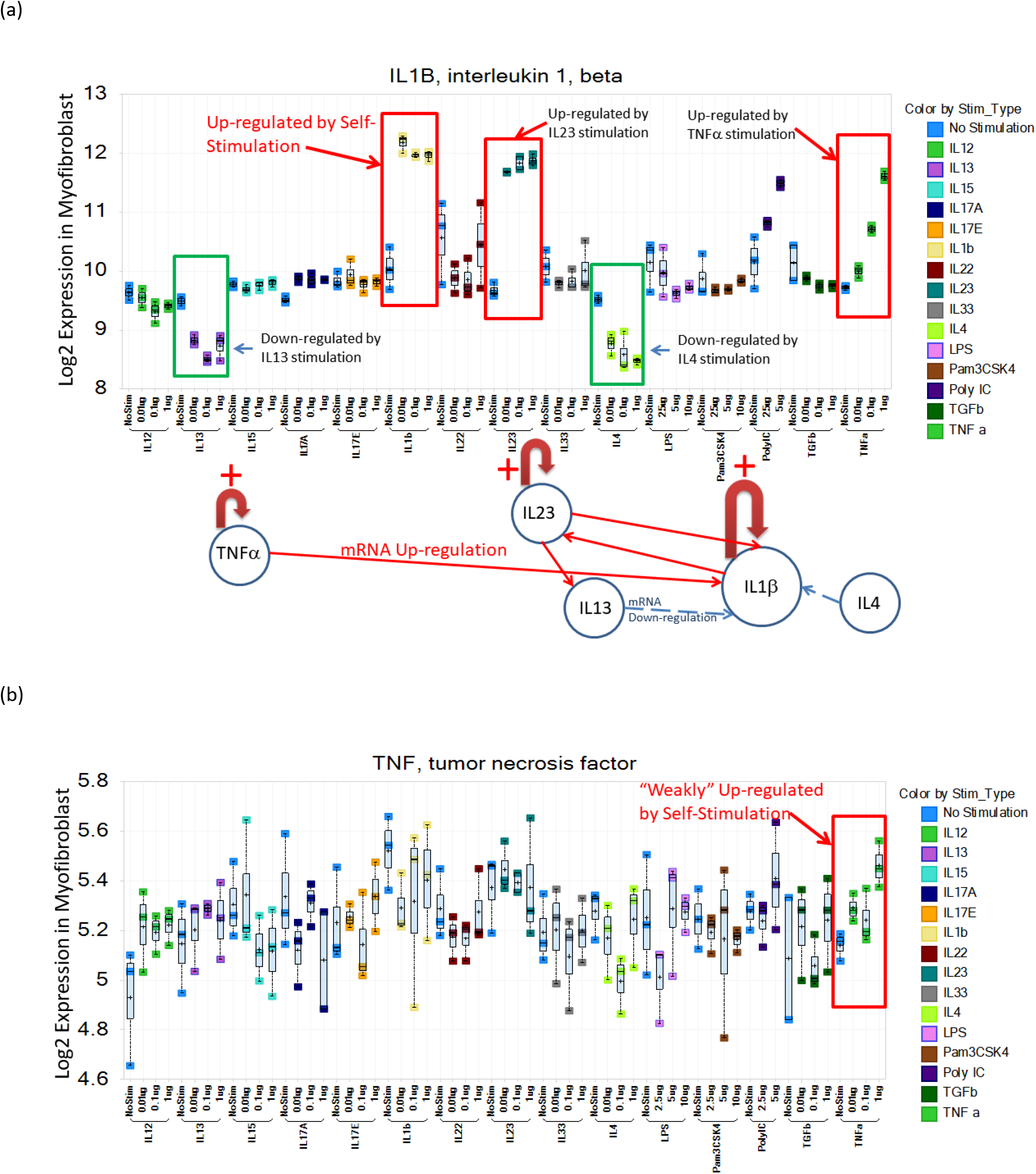
Gene expression profiles of IL-1β (a) and TNFα (b) in the Intestinal Myofibroblast cells under 15 stimulation conditions presented by the color scheme on the right. Same as in Figure 2, stimulation conditions that lead to significant changes of IL-1β or TNFα expressions are marked by square boxes where at least one dose of a specific cytokine stimulation is significantly different from its associated non-stimulated control samples (blue dots). The network plot between panels summarizes the interactions between 5 cytokines where red lines represent up-regulate its target expression and dashed lines mean down-regulation. Amplification by self-stimulation of TNFα, IL-1β and IL-23 are noted.

Figure 5 summarizes the cytokine regulatory networks from the intestinal primary myofibroblast cell stimulation experiment where IL-1β, IL-23 and TGFβ1 were found to be sensitive to self-stimulation (marked in red). This is very different from the cytokine networks shown in Figure 3 for HCT8 cells where TNFα was the only “causal” node identified. Even though TNFα still appears to be a central node controlling several other cytokines and was up-regulated by self-stimulation slightly (~1.2fold change), it did not reach the self-regulation threshold of 1.5-fold. This may suggest that in the myofibroblast, there is less potential TNFα self-amplification. Other genes and proteins still respond to TNFα, but myofibroblast cells themselves might not be a robust source of TNFα to sustain the downstream effects of this cytokine. On the other hand, IL-1β, IL-23 and TFGβ1 appear to be self-amplifying and likely to be sensitive and “causal” in the cytokine network. In summary, in myofibroblast, blocking or decreasing the levels of IL-1β, IL-23 or TGFβ1 would likely have a stronger effect on downstream gene expression.

**Figure 5.**
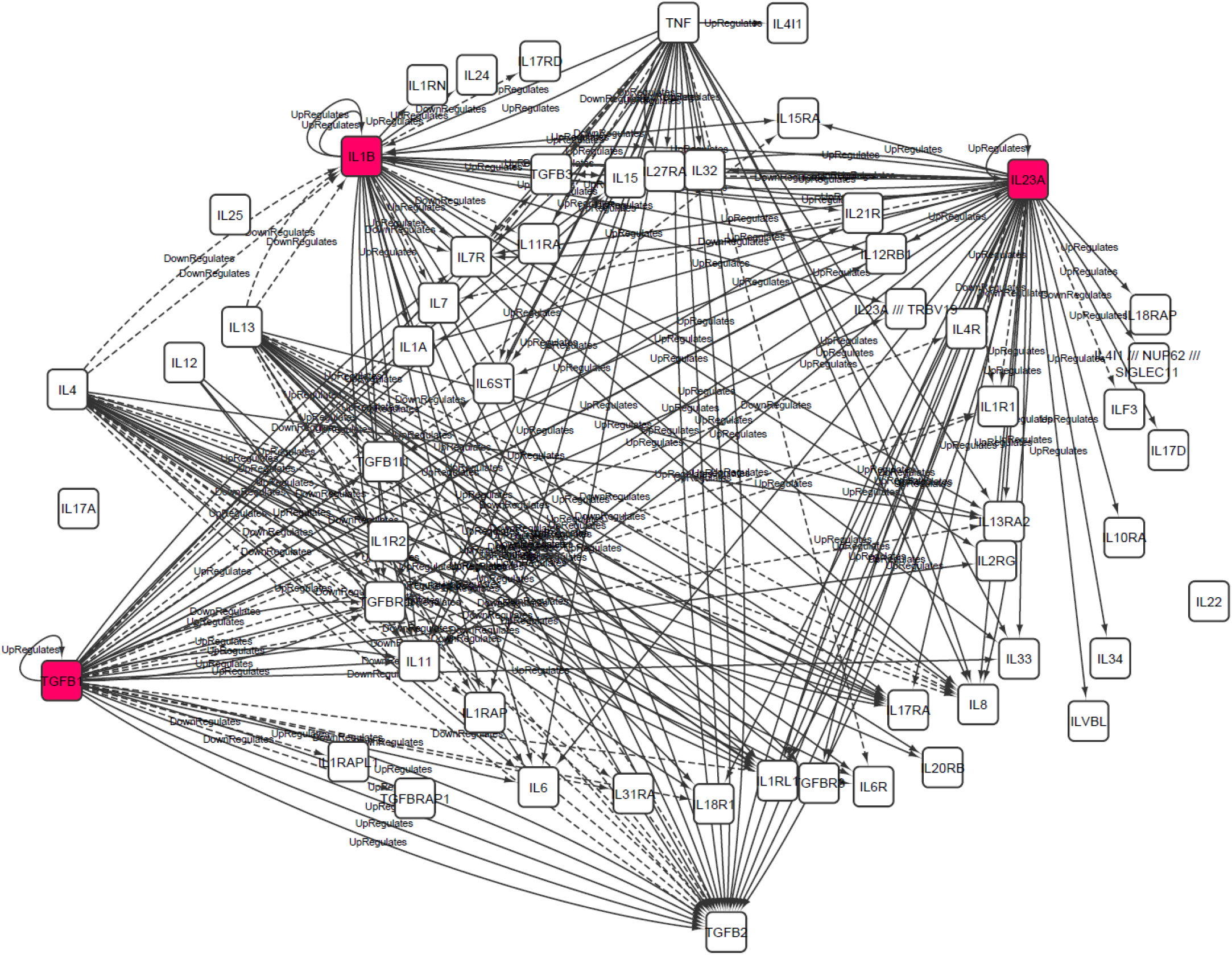
Cytokine network based upon the stimulation experiment from intestinal primary myofibroblast using 15 stimulation conditions described in Figure 4. Same criteria used as in Figure 3 – only cytokine nodes are plotted here, solid lines represent up-regulation and increasing expression of target cytokines, and dashed lines represent down-regulation. Here IL-1β, IL23 and TGFβ1 have self-loops because they were up-regulated by self-stimulation and shown as red boxes.

## IV. Discussion: Criteria to support or refute the idea – literature review for existing cytokine targets

Colon epithelial cell modeled by the HCT8 cell culture system is an important tissue type involved in Inflammatory Bowel Disease (see Parikh et al. 2019), and we know that TNFα is significantly up-regulated in colon samples from IBD patient as compared to healthy human subjects (Olsen et al. 2007). TNFα’s self-positive regulation in HCT8 cell lines would suggest it might play a “causal” role in the IBD disease phenotype (Roulis et al. 2011), at least for certain patient populations. It is noted here that a single cultured cell type is not enough to describe the complex environment of disease tissue with many cell types, so identifying positive feedback loops of TNFα in Crohn’s (CD) or Ulcerative Colitis (UC) disease tissues would more directly support the disease causal functions of TNFα in IBD. Performing cytokine perturbation experiments on the whole diseased tissue to identify if there exist self-positive regulations of a specific target will reveal more disease specific cytokine networks. For many diseases, inflamed tissues are difficult to obtain thus perturbation experiments on the disease tissues are not easy to perform, thus data from such experiments would be very valuable for improving our understandings of molecular interaction networks that specific for a disease mechanism.

As mentioned above, even though cytokine interaction network models from direct perturbation experiments on disease tissues are very limited at present time, many existing publications provided some indirect evidence or non-quantitative models of positive-feed-loop of several cytokines and their pathological roles in several inflammatory diseases. Below we briefly review such literature support in different auto-inflammatory diseases.

1. **Crohn’s Disease and Ulcerative Colitis (CD/UC)** – Beyond evidence for TNFα’s “positive-self-regulation” from the HCT8 cell stimulation experiment, a previous PNAS paper (Roulis et al. 2011) demonstrated overproductions of TNFα in intestinal epithelial cell (IEC) play “causal” role in IBD by showing “chronic overproduction of TNFα by IEC suffices to cause full development of Crohn-like pathology” in a mouse model. Notice that “chronic overproduction of TNFα” in the intestinal epithelial cell is generated by a “self-positive feedback loop” of TNFα after a genetic mutation of TNF regulatory element leading to its self-sustained over-production (Kontoyiannis et al. 1999). Such phenomenon clearly supports the idea that positive-feedback loops of TNFα could be causal for UC and CD.
2. **Psoriasis/ Psoriatic Arthritis/ Ankylosing Spondylitis** – – majority studies for these three diseases are concentrated on the up-regulation of relevant cytokines in the disease but studies of their positive-feedback mechanism are emerging, especially for Psoriasis (Lowes et al, 2013; Towne & Sims, 2012). There are growing clinical evidence to demonstrate that IL-23, IL-17 and TNFα are effective targets for Psoriasis with different degrees of efficacy. Many research reports and literatures support the idea of cytokine network including IL-23, IL-17A/F/C, TNFα, and additional cytokines such as IL-36G are involved in the Psoriasis pathogenesis (reviewed by Lowes et al. 2013), researchers are beginning to shift from a pathway-centered view into a more network-based view based on multiple feedback loops identified in Psoriasis (Lowes et al., 2013). Results and data accumulated from the clinical and basic research of Psoriasis thus are beginning to reveal the interesting cytokine network involved in Psoriasis pathogenesis, and how important cytokine nodes such as IL-23, IL-17 and TNFα in the Psoriasis disease network behave in the disease progression and treatment responses based on targeting those cytokines. Revealing the cytokine networks involved in Psoriasis might be useful for the disease mechanism understandings of other autoimmune diseases.
3. **Rheumatoid Arthritis (RA)**: Anti-TNFα therapy has been effective for RA, and this agrees with the “Self-Positive Regulation” of TNFα in RA supported by data presented by Lai et al. (2014). IL-6 as an effective RA target is supported by Mori et al. 2011 reporting “STAT3-dependent cytokine amplification in RA”, notice Figure 7 in the paper specifically plotted (positive) feedback loop involving IL-6, which suggests existence of IL-6 “self-positive regulation” and its “causal” role in RA. Another supporting data for targeting IL-6 in RA is suggested by Das et al. (2014) where they reported patients who treated with anti-IL6R therapy tocilizumab had better clinical responses if they had higher serum IL-6 levels on the baseline. This suggests that from biomarkers perspective targeting IL-6 is clinically effective.
4. **Giant Cell Arteritis (GCA)** – there are conflicting reports about whether TNFα, IL1b, and IL-6 play “causal” role in GCA and also whether those targeted therapies are truly effective. Even though their expression levels were elevated in the relevant tissue (Hernandez-Rodriguez et al., 2004), clinical studies (Hoffman GS, Cid MC, Weyand CM, et al., 2005; Visvanathan et al., 2011) suggest that “although TNF is found in abundance in affected vessels, it might not play a critical role in the pathogenesis of GCA”. Overall, current literatures suggest the correlations of above cytokines with GCA, but the “causality” issue is not determined and hence the efficacy of any of those targeted therapies, same with IL-17 and IL-12/23 related approaches (Furst & Emery, 2014). All these evidence raise the importance of identifying “causal” cytokines in this disease, which could be potentially answered by the approach advocated in this paper. More data are needed to show whether identifying “positive-self regulation” could be useful to identify a more “causal” cytokines for GCA, and to support or reject the idea described here.
5. **Juvenile Idiopathic Arthritis (JID/AID)** – JID is complex and heterogeneous with at least five subtypes (Mangge & Schauenstein, 1998), and literatures have suggested the importance of “positive and negative cytokine feedback” in this disease (Woo, 2002). Anti-TNFα and Anti-IL-6 therapies have shown some effectiveness for JIA (Semeraro et al., 2014), but evidence of “positive-self-regulation” are not studied beyond up-regulation in the diseases state. There is one paper (Dinarello 2005) describing effectiveness of anti-IL-1β in systemic onset juvenile idiopathic arthritis (SoJIA), it specially pointed out in the abstract “**Loss of control of the secretion of IL-1β might be a common mechanism explaining the aberrant activity of IL-1 in these diseases**", thus confirming the out of control “positive-regulation” of IL-1β in this disease type and suggesting its causal role.
6. **Gout** – gout is induced by inflammatory interactions between monosodium urate (MSU) crystals and local tissues (Dalbeth & Haskard, 2005). A recent paper (Mylona et al, 2012) described the increased production of IL-1β in Gout: “MSU crystals alone failed to induce IL-1β, IL-6 or TNFα in both patients and control groups, but a stronger synergy between MSU/Pam3Cys and MSU/C18:0 for the **induction of IL-1β** was found in patients with gout compared to healthy controls.… The synergy between MSU crystals and TLR-2 ligands is more prominent in patients with gout than in controls. This is likely mediated by the **enhanced maturation of pro-IL-1β into IL-1β**”. The first sentence quoted here indicated that IL-6 and TNFα are not induced in gout thus suggests they are not likely to be good targets. Second sentence quoted shows IL-1β is up-regulated and possibly by a self-positive loop (i.e., converting pro-IL-1β to IL-1β), thus supporting the idea of “positive-self-regulation” of IL-1β in Gout and the effectiveness of anti-IL-1β for this disease.
7. **Multiple Sclerosis** – the failure of anti-TNFα therapy for MS could be a clear refuting case for the idea described here because TNFα levels are shown to be elevated in MS patients (e.g., Maimone et al, 1991; Hohnoki et al., 1998) no withstanding whether it is “positively-self-regulated” or not. One explanation proposed by Robinson et al. (2001), is related to the blood-brain barrier which prevents the anti-TNFα agent’s entry to the CNS. That effect is assumed to lead to an increase in peripheral T cell auto-reactivity. This explanation thus again emphasizes the importance of homeostatic levels of TNFα both in the peripheral and in the CNS in relation to the MS associated symptom, thus the importance of any potential “positive” or “negative” self-regulation of TNFα in this CNS disease.

## V. Summary and Conclusion

In this paper a potential method to test “causality” in a cytokine network is proposed, some preliminary data analyses and literature evidence supporting the relevance of “positive-feedback” to a cytokine targets’ causal role for disease are also listed. Below we discuss the implications of this method and its some potential utilities and next step.

1. **Positive Self-Regulated Cytokines causal for diseases – Dynamic***cis***-acting elements?**-- Causality idea presented here is similar to the gene network approach using QTL and Bayesian network approach (Chen et al., 2008) but extend to cytokines/genes that might not directly under genetic control or lacking SNPs and genetic perturbation information. For immune systems that operate in hours to days, extracting this type of “dynamical” causal interactions based upon feedback self-regulations might provide some important complimentary novel information not captured by the genetics approach. The “Positive Self-Regulation” concept is similar to the idea of “*cis*-acting” element or “*cis*-eQTLs” but without the genetic variations, in the sense that the causality here means a gene/protein (positively) controls and amplifying its own levels or even leads to an out-of-control/ dis-regulated level thus ultimately controls other genes and disease phenotypes dynamically.
2. **Single cell type or whole tissue?** – It should be noted that data presented in this paper are from two individual cell types, which might be only a small part of any complex tissue type in a healthy or diseased state. To identify the most important causal cytokine in an inflammatory disease, re-constructing the cytokine interactions in the whole inflamed tissue will be very important and crucial since that is the most relevant state, but of course such experiments are more difficult to perform because a lot of diseased tissues are not easy to obtain. Whether a whole tissue cytokine interaction network can be estimated from networks derived from individual cell types will be an interesting question to test.
3. **Extend to targets other than cytokines** – Idea proposed here can be used to select/differentiate other type of disease targets, such as kinases (PI3K, JAKs), GPCRs, co-signaling molecules etc. It is likely that there may be more than one “causal” cytokines/proteins that could be important for a specific disease, this causal information can be used to potentially select a specific kinase target/compound based upon whether it impacts the desired subsets of cytokines/proteins that were shown being causal for the disease phenotype.
4. **Immune-regulations by T-regs and beneficial bacterial species** – In the above we concentrated on identifying causal cytokines that are often increased or up-regulated in a diseased state and are mostly pro-inflammatory in nature. We also know that some cytokines (such as IL-4, IL-13 and TGFβ1 shown in the myofibroblast cells) can be anti-inflammatory in certain conditions and down-regulate pro-inflammatory cytokines. Targeting those anti-inflammatory cytokines/pathways is clearly an important disease-interception strategy. One possible function of anti-inflammatory cytokines and beneficial bacterial species in the relevant tissue environment may be their abilities to *stabilize* the pro-inflammatory cytokines that are in a dis-regulated and “causal” state for disease. Utilizing body’s intrinsic anti-inflammatory mechanism to achieve an immune balance and reduce inflammation will be a smarter way of control positive feedback loop in auto-immune diseases for future research.

In conclusion, here a cytokine network idea is proposed to test the dynamic “causality” of disease-relevant cytokines/proteins, and such methods could be useful for future target selection/differentiation for auto-immune diseases. As a target selection and disease interception strategy, we start with considering cytokines that are more expressed in disease than in normal healthy state just like most current research approaches. Many those up-regulated genes are selected as drug targets for certain diseases based upon literature and biological supports for their disease association. Idea described here then could be used to differentiate whether an up-regulated gene/protein is “causal” for a disease or just only correlates with the disease phenotype and might be controlled by more important genes/proteins. Differentiating “causal” versus “reactive” nodes in the cytokine networks using the method proposed here could improve our understanding of disease mechanisms and hopefully lead to discoveries of better medicine in the future.

## Acknowledgement

This paper is based upon an abstract and poster presented at Keystone Symposium on System Immunology (Chen et al. 2015). The author would like to acknowledge Mamatha Challapalli for her experimental expertise and Drs. Marcos Milla, Xuejun Liu and Oivin Guicherit for valuable discussions and editing help.

## References

Caldwell, AB, Cheng, Z, Vargas, JD, Birnbaum, HA, & Hoggmann, A (2014). Network dynamics determine the autocrine and paracrine signaling functions of TNF. Genes & Development. 28:2120–2133. doi:10.1101/gad.244749.114.

Chen, Y, Challapalli, M, Liu, X, Guicherit, O, Milla, M (2015) A Network View of Cytokine Interaction: Method to construct “Causal Cytokine Networks” for Disease Target Identification and Differentiation. Keystone Symposium on System Immunology Abstract and Poster.

Chen, Y*, Zhu, J*, Lum, PY, Yang, X, Pinto, S, MacNeil, DJ, Zhang C, Lamb, J, Edwards, S, Sieberts, SK, Leonardson, A, Castellini, LW, Wang, S, Champy, M-F, Zhang B, Emilsson, V, Doss S, Ghazalpour A, Horvath S, Drake, TA, Lusis, AJ, Schadt, EE (2008). Variations in DNA elucidate molecular networks that cause disease. Nature, 452:429–435. (*Equal Contribution).

Cohen, J D, Bournerias I, Buffard V et al. 2007. Psoriasis Induced by Tumor Necrosis Factor-alpha Antagonist Therapy: A Case Series. J Rheumatol 2007;34:380–5. https://www.jrheum.org/content/34/2/380.long

Dalbeth, N. & Haskard D.O. 2005. Mechanisms of inflammation in gout. Rheumatology 2005;44:1090–1096 doi:10.1093/rheumatology/keh640

Das S, Vital EM, Harton S, Bryer D, El-Sherbiny Y, Rawstron AC, Ponchel F, Emery P, Buch MH. 2014 Abatacept or tocilizumab after rituximab in rheumatoid arthritis? An exploratory study suggests non-response to rituximab is associated with persistently high IL-6 and better clinical response to IL-6 blocking therapy. Ann Rheum Dis 2014;73:909–912. doi:10.1136/annrheumdis-2013-204417

Dinarello, C.A. 2005, Blocking IL-1 in systemic inflammation, J Exp Med. 2005 May 2;201(9):1355–9.

Farahnik, B., Beroukhim, K., Nakamura,M., et. al. (2016) Anti-IL-17 Agents for Psoriasis: A Review of Phase III Data. J. Drugs Dermatol. 15(3):311–316

Furst D E, & Emery P, 2014. Rheumatoid arthritis pathophysiology: update on emerging cytokine and cytokine-associated cell targets. Rheumatology (Oxford). 2014 Jan 8. [Epub ahead of print]

Hernandez-Rodriguez J, Segarra M, Vilardell C. et al. 2004. Tissue production of pro-inflammatory cytokines (IL-1b, TNFa and IL-6) correlates with the intensity of the systemic inflammatory response and with corticosteroid requirements in giant-cell arteritis. Rheumatology, 43:294–301 doi:10.1093/rheumatology/keh058.

Hoffman GS, Cid MC, Weyand CM, et al. Phase II study of the safety and efficacy of infliximab in giant cell arteritis (GCA): 22 Week interim analysis. Arthritis Rheum 2005; 52(suppl):S271.

Hohnoki K, Inoue A, Koh CS. 1998. Elevated serum levels of IFN-gamma, IL-4 and TNF-alpha/unelevated serum levels of IL-10 in patients with demyelinating diseases during the acute stage. J Neuroimmunol. 1998 Jul 1;87(1-2):27–32

Kontoyiannis D, Pasparakis M, Pizarro TT, Cominelli F, Kollias G (1999) Impaired on/off regulation of TNF biosynthesis in mice lacking TNF AU-rich elements: Implications for joint and gut-associated immunopathologies. Immunity 10:387–398.

Lai Y, Bai X, Zhao Y et al., ADAMTS-7 forms a positive feedback loop with TNF-α in the pathogenesis of osteoarthritis. Ann Rheum Dis. 2014 Aug;73(8):1575–84. doi: 10.1136/annrheumdis-2013-203561. Epub 2013 Aug 8.

Jakobosen M et al. 2009. Amelioration of Psoriasis by Anti-TNF-α RNAi in the Xenograft Transplantation Model. Molecular Therapy (2009) 17(10), 1743–1753. doi:10.1038/mt.2009.141

Konsta M, Bamias G, Tektonidou M G, et al. 2013. Increased levels of soluble TNF-like cytokine 1A in ankylosing spondylitis. Rheumatology (2013) 52 (3): 448–451. doi: 10.1093/rheumatology/kes316

Maimone D, Gregory S, Arnason BG, Reder AT. 1991. Cytokine levels in the cerebrospinal fluid and serum of patients with multiple sclerosis. J Neuroimmunol. 1991 Apr;32(1):67–74.

Mangge H, & Schauenstein K, 1998, Cytokines in juvenile rheumatoid arthritis (JRA). CYTOKINE, 10(6), June, 1998:471–480

Morel PA, Lee REC, Faeder JR. 2017. Demystifying the cytokine network: Mathematical models point the way. Cytokine, 98:115–123.

Mori T, Miyamoto T, Yoshida H, Asakawa M, Kawasumi M, Kobayashi T, Morioka H, Chiba K, Toyama Y, Yoshimura A 2011. IL-1β and TNFα-initiated IL-6–STAT3 pathway is critical in mediating inflammatory cytokines and RANKL expression in inflammatory arthritis, International Immunology, Volume 23, Issue 11, November 2011, Pages 701–712, https://doi.org/10.1093/intimm/dxr077

Mylona EE, Mouktaroudi M, Crisan TO, et al. 2012. Enhanced interleukin-1β production of PBMCs from patients with gout after stimulation with Toll-like receptor-2 ligands and urate crystals. Arthritis Research & Therapy 2012, 14:R158 doi:10.1186/ar3898

Olsen T, Rasmus Goll R, Cui G, Husebekk A, Vonen B, Birketvedt G, & Florholmen J (2007) Tissue levels of tumor necrosis factor-alpha correlates with grade of inflammation in untreated ulcerative colitis, Scandinavian Journal of Gastroenterology, 42:11, 1312–1320, DOI: 10.1080/00365520701409035

Parikh K, Antanaviciute A, Fawkner-Corbett D, Jagielowicz M, Aulicino A, Lagerholm C, Davis S, Kinchen J, Chen HH, Alham NK, Ashley N, Johnson E, Hublitz P, Bao L, Lukomska J, Andev RS, Björklund E, Kessler BM, Fischer R, Goldin R, Koohy H, Simmons A. Colonic epithelial cell diversity in health and inflammatory bowel disease. Nature. 2019 Mar;567(7746):49–55. doi: 10.1038/s41586-019-0992-y. Epub 2019 Feb 27. PMID: 30814735.

Tobin, A M, Kirby B. 2005. TNF alpha inhibitors in the treatment of psoriasis and psoriatic arthritis. BioDrugs. 2005;19(1):47–57.

Robinson W H, Genovese M C, Moreland L W. 2001. Demyelinating and neurologic events reported in association with tumor necrosis factor alpha antagonism: by what mechanisms could tumor necrosis factor alpha antagonists improve rheumatoid arthritis but exacerbate multiple sclerosis? Arthritis Rheum. 2001 Sep;44(9):1977–83.

Roulis M, Armaka M, Manoloukos M, Apostolaki M, and Kollias G, 2011. Intestinal epithelial cells as producers but not targets of chronic TNF suffice to cause murine Crohn-like pathology. PNAS 2011 108 (13) 5396–5401; published ahead of print March 14, 2011, doi:10.1073/pnas.1007811108

Schett G, Elewaut D, McInnes I B, Dayer J-M, & Neurath M F. 2013. How Cytokine Networks Fuel Inflammation: Toward a cytokine-based disease taxonomy. Nature Medicine, (19)822–824 DOI: doi:10.1038/nm.3260.

Semeraro F, Arcidiacono B, Nascimbeni G, Angi M, Parolini B, Costagliola C. 2014, Anti-TNF therapy for juvenile idiopathic arthritis-related uveitis. Drug Design, Development and Therapy, March 2014 Volume 2014:8 Pages 341–348, DOI: http://dx.doi.org/10.2147/DDDT.S54207

Visvanathan S, Rahman MU, Hoffman GS, Xu S, García-Martínez A, Segarra M, Lozano E, Espígol-Frigolé G, Hernández-Rodríguez J, Cid MC. 2011. Tissue and serum markers of inflammation during the follow-up of patients with giant-cell arteritis--a prospective longitudinal study. Rheumatology (Oxford). 2011 Nov;50(11):2061–70. doi:10.1093/rheumatology/ker163. Epub 2011 Aug 25.

Waito M, Walsh SR, Rasiuk A, Bridle BW, Willms AR 2016. A Mathematical Model of Cytokine Dynamics During a Cytokine Storm. In: Bélair J., Frigaard I., Kunze H., Makarov R., Melnik R., Spiteri R. (eds) Mathematical and Computational Approaches in Advancing Modern Science and Engineering. Springer, Cham. https://doi.org/10.1007/978-3-319-30379-6_31

Woo, P. 2002, Cytokines and juvenile idiopathic arthritis. Curr Rheumatol Rep. Dec;4(6):452–7.

Yiu HH, Graham AL, Stengel RF 2012. Dynamics of a Cytokine Storm. PLoS ONE 7(10): e45027. https://doi.org/10.1371/journal.pone.0045027

